# Inexpensive Multiplexed Library Preparation for Megabase-Sized Genomes

**DOI:** 10.1101/013771

**Authors:** Michael Baym, Sergey Kryazhimskiy, Tami D. Lieberman, Hattie Chung, Michael M. Desai, Roy Kishony

**Affiliations:** Department of Systems Biology, Harvard Medical School, USA; Department of Organismic and Evolutionary Biology; FAS Center for Systems Biology; Department of Physics, Harvard University, USA; Faculty of Biology and Life Sciences & Engineering, Technion, Israel.

**Author notes:** These authors contributed equally to this work (Michael Baym, Sergey Kryazhimskiy, Tami D. Lieberman, Hattie Chung).

## Abstract

Whole-genome sequencing has become an indispensible tool of modern biology. However, the cost of sample preparation relative to the cost of sequencing remains high, especially for small genomes where the former is dominant. Here we present a protocol for the rapid and inexpensive preparation of hundreds of multiplexed genomic libraries for Illumina sequencing. By carrying out the Nextera tagmentation reaction in small volumes, replacing costly reagents with cheaper equivalents, and omitting unnecessary steps, we achieve a cost of library preparation of $8 per sample, approximately 6 times cheaper than the widely-used Nextera XT protocol. Furthermore, our procedure takes less than 5 hours for 96 samples and uses nanograms of genomic DNA. Many hundreds of samples can then be pooled on the same HiSeq lane via custom barcodes. Our method is especially useful for re-sequencing of large numbers of full microbial or viral genomes, including those from evolution experiments, genetic screens, and environmental samples.

## Introduction

Sequencing has become an indispensible tool in modern microbiology, dramatically changing the resolution and speed of studies of biodiversity [1], evolution [2-6], and molecular biology [7], and improving pathogen surveillance [8] and clinical diagnostics [9,10]. With current technology, hundreds of full megabase-size genomes can be sequenced in a single Illumina HiSeq lane at over 30x coverage, for a cost of about $15 per sample. Thus, the costs of standard library preparation methods, which typically exceed $50 per sample, substantially limit the amount of microbial genome sequencing. Two studies have recently proposed ways to alleviate this limitation [11,12]. Based on similar principles to those proposed by Lamble et al. [12] and in the Illumina Nextera XT kit [13], we developed a library-preparation protocol that achieves further reductions in costs and increases in efficiency. The protocol described here costs approximately $750 per 96 samples including consumables, with under 3 hours hands on time and under 5 hours total time.

## Protocol overview and important considerations

Our protocol consists of 5 modules (Figure 1). We assume that the protocol is executed with purified genomic DNA (gDNA) but other types of purified DNA can be used. Since the reliability of the tagmentation reaction (Module 2) is sensitive to the purity of input gDNA [12], we recommend using column-based genomic extraction, such as the Invitrogen PureLink 96-well kit. The cost of consumables per sample is rounded to the nearest $0.25.

**Figure 1.**
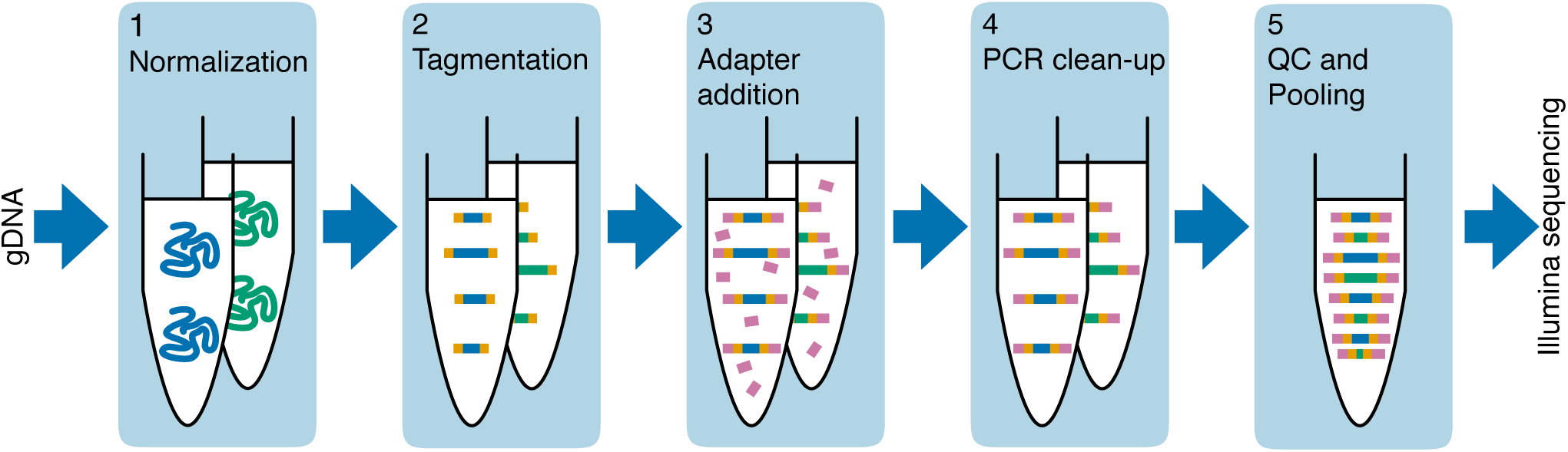
Schematic of library preparation workflow.

### Module 1: Normalization of gDNA concentrations across samples ($0.50/sample, 60 min)

The goal of this module is to normalize the gDNA concentration across samples to achieve uniform reaction efficiency in the tagmentation step (Module 2). Tagmentation is sensitive to the input gDNA concentration and the optimal concentration will vary depending on the organism, DNA type (e.g., genomic versus PCR product), and the DNA extraction method. See “Selecting input gDNA concentration and bead volume for optimal fragment length” and Figure 2 for more information. In general, we found that a starting concentration of 0.5ng/μl works well for gram-negative bacteria.

**Figure 2.**
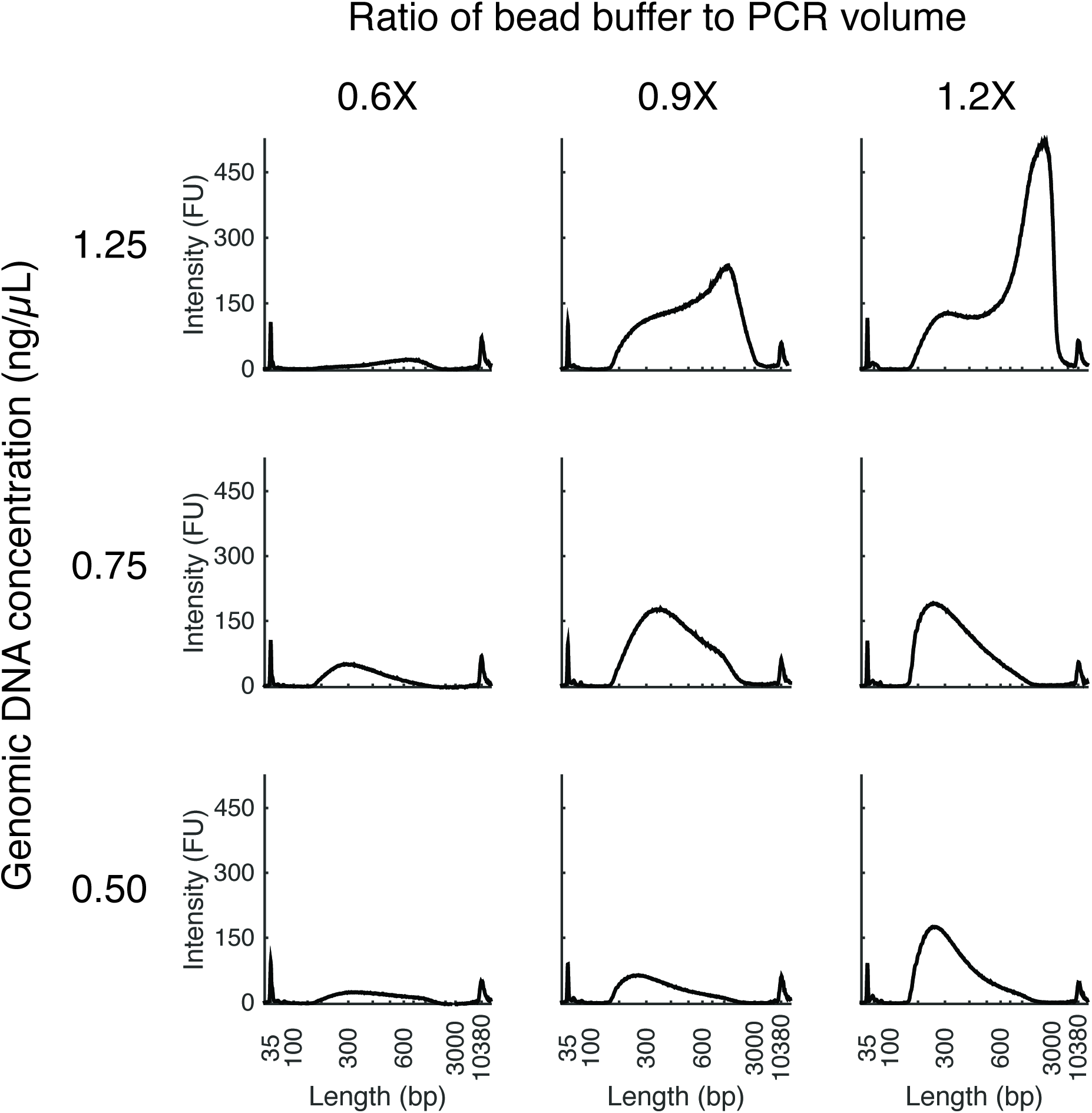
Dependence of fragment size distribution on input gDNA concentration and bead volume. DNA fragment size distribution is affected by starting genomic DNA concentration (rows) as described in Module 1 as well as the relative amount of bead buffer used in PCR clean-up (columns) as described in Module 4. Size distribution is measured by BioAnalyzer and reported in fluorescence units. Data is from *Stenotrophomonas maltophilia* (67% GC, 4.8 Mb genome). At high initial gDNA concentration (1.25 ng/μl) the fragment distribution is right-skewed, suggesting the presence of chimeras.

We use SYBR Green I to quantify gDNA, which gives sufficiently precise measurements and is markedly cheaper than other dyes. For lower-throughput work QuBit quantification can also be used. We do not recommend absorbance quantification methods such as NanoDrop because they have lower sensitivity and can be affected by the presence of single-stranded nucleic acids.

### Module 2: Tagmentation ($4.75/sample, 30 min)

In this module, the transposase loaded with Illumina adaptors (also referred to as “tagmentation enzyme”) and the tagmentation buffer provided in an Illumina Nextera kit are used to simultaneously fragment gDNA and ligate sequencing adaptors. We use steps described in the standard Nextera protocol, but with a smaller reaction volume. We found that tagmentation-reaction volume as small as 2.5μl does not significantly limit the diversity of sequenced DNA for megabase-sized genomes. Specifically, libraries of *E. coli* (genome size 4.64Mb) prepared with this protocol typically yield about 97% unique reads. Since each position in the genome is represented on thousands of tagmented DNA fragments, the fraction of false-positive variants created by errors in subsequent PCR amplification (Module 3) will be negligible. Larger tagmentation-reaction volumes may be necessary for larger genomes to achieve sufficient library complexity and avoid PCR-induced errors. The final fragment size distribution critically depends on the stoichiometry of gDNA and transposase [14]. Thus, to achieve results consistent across samples, it is essential to accurately normalize input DNA (Module 1) and to thoroughly mix the tagmentation master mix. On the other hand, purification after tagmentation is unnecessary, and tagmented DNA can be directly used as template for the subsequent PCR step.

### Module 3: Adapter addition and library amplification ($1.50/sample, 75 min)

In this module, PCR is used to attach Illumina adaptors and sample barcodes to tagmented DNA fragments. The adaptors bind fragments to the flow cell [14], and the barcodes allow for multiplexed sequencing. If 96 or fewer samples are pooled, we use the Illumina TruSeq primers S501-S508 and N701-N712. If higher multiplexing is required, we developed custom row and column primers, labeled R09-R36 and C13-C24. These were derived from the TruSeq primers and are compatible with them (Supplementary File 1).

The number of PCR steps and the PCR volume may be adjusted, if necessary. However, with a large number of PCR cycles chimera formation may become a problem [15]. These chimeras do not affect the Illumina sequencing reaction because single-stranded DNA fragments attach to the flow cell [16]. They can however strongly bias the fragment-size estimation, which decreases the number and quality of sequence reads (see Section “Selecting input DNA concentration and bead to sample ratio”). A peak at over 1kb in the BioAnalyzer trace of the library often indicates excessive chimera presence. While it is possible to successfully sequence despite chimera presence (see Figure 3), should chimeras be a problem we recommend either a “reconditioning PCR”, i.e., 3-4 additional cycles with fresh primers and polymerase [15], or a higher initial primer concentration.

**Figure 3.**
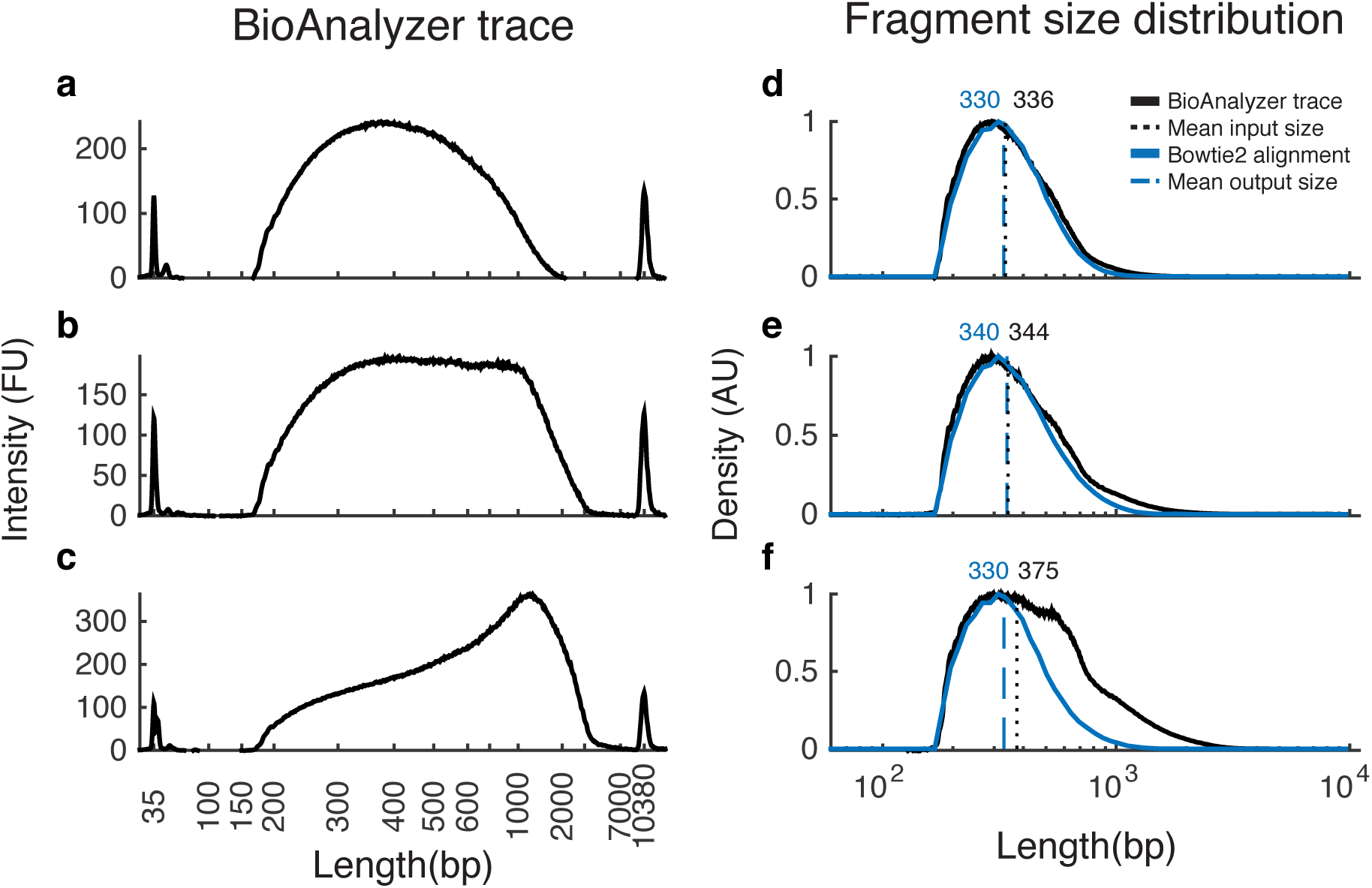
Correspondence between BioAnalyzer traces and the length distribution of aligned reads. Panels A-C show three representative BioAnalyzer traces. Panels D-F show the corresponding estimated fragment-size distribution (black) and distribution of sequenced reads calculated from aligning to a reference genome (blue). We estimate the relative abundance of fragments from a BioAnalyzer trace by dividing the relative fluorescence intensity by the fragment length and rescale the histogram to the change of axes. Note that sequencing can be successful despite the presence of apparently very long fragments (which are likely chimeras) in the BioAnalyzer traces (Panels C and F). All data is from *S. maltophilia*.

### Module 4: PCR clean-up and size selection ($0.50/sample, 40 min)

In this module, PCR products are purified with magnetic beads and enriched for fragments of the desired length for sequencing. In lieu of the significantly more expensive Illumina-recommended AmPure XP beads, we use magnetic bead solution with the “MagNA” bead-extraction protocol from [11,17] and Thermo Sera-Mag SpeedBeads. See the “Selecting input DNA concentration and bead to sample ratio” section for more information, and Figure 2 for an example. In general, we found that a 1:1 volumetric ratio of sample to beads works well.

### Module 5: Library quality control and pooling ($1/sample, 90 min)

In this module, sample concentrations and fragment size distributions are estimated and libraries are pooled. We measure the DNA concentration of each sample fluorescently, as in Module 1; quantification by qPCR is unnecessary at this stage. We discard samples with less than 0.5ng of DNA. Fragment size distribution can be measured with an Agilent BioAnalyzer or TapeStation. While it would be ideal to measure the size distribution of every sample, this is not practical or economically feasible at large scale. Moreover, we found that sample preps from the same 96-well plate typically have similar post-cleanup fragment-size distributions. Thus, we estimate this distribution for a subset of samples (5 to 10). Then, based on individual sample concentrations and the common average fragment length, we calculate the DNA molarity of each sample and pool variable volumes of samples to achieve equimolar concentrations in the pool. A final verification of the pooled sample concentration can be useful before sequencing (including qPCR), though most sequencing centers perform this as part of sample quality control.

### Selecting input DNA concentration and bead to sample ratio

For each sample, this protocol produces an adaptor-ligated, barcoded, library of DNA fragments with some distribution of read lengths. The optimal fragment-size distribution depends on the sequencing protocol. To maximize the amount of useful sequenced DNA, most fragments should be longer than the combined length of the sequencing reads and adaptors (e.g. above 338bp for paired-end 100bp sequencing). However, as longer fragments are underrepresented in aligned sequencing reads (Figure 3) and can lower overall cluster density (Illumina Nextera technical notes), fragment length should ideally be kept under 1kb.

We found that the distribution of fragment sizes depends strongly on two parameters: (1) the input concentration of gDNA during tagmentation (output of Module 1) and (2) the amount of beads during post-PCR clean-up (Module 4). Higher input gDNA concentrations generally result in longer fragments because each tagmentation enzyme makes a single cut [13]. Higher bead to sample ratios for purification can yield smaller fragments because beads preferentially capture larger fragments[17]. We recommend calibrating these two parameters on a representative sample prior to scaling up the preparation. For example, the initial concentration of DNA might be increased to increase average fragment size.

We tested this protocol exclusively with Illumina sequencing. However, by changing the barcode oligos, it can be modified for any adaptor-based sequencing platform (e.g., Ion Torrent, 454, SOLiD, and PacBio [13]).

## Detailed Protocol

This protocol is for the preparation of 96 samples (8 rows x 12 columns) but can be modified for either higher or lower throughput.

### General tips

- We advise cleaning pipettes and your station with DNA-Away (Thermo Scientific 7010) to reduce contamination from the environment and previous samples.
- Many steps call for centrifugation of 96-well plates. These steps are necessary for consistent multi-channel small volume liquid transfer and should not be omitted.
- The potential for cross-contamination of samples, or particularly primers, is high. We recommend the use of filter tips, exclusively aspirating samples or primers with fresh tips, and avoiding “blowing out” the pipette.

### Materials and equipment used throughout the protocol

- Sterile DNase-free water
- Filter tips
- Manual multi-channel pipettes. We found less consistent results when mixing with electronic multi-channel pipettes.
- DNase-free microfuge tubes, PCR strips, PCR tubes, and PCR 96-well plates
- Centrifuge capable of spinning 96-well plates
- Thermocycler
- (optional) Electronic multi-dispense, multi-channel pipettes
- (optional) 96-well pipette (Liquidator or equivalent)
- (optional) PCR cooler (e.g. Eppendorf 3881)
- (optional) Rubber roller for sealing plates
- (optional) Liquid-handling robot

## Module 1. DNA normalization

**Goal:** Obtain at least 5μl of each sample at concentration 0.5ng/μl (or another chosen concentration)

### Materials and equipment

- gDNA in 96-well plate in range of about 1-10ng/μl
- TE buffer
- 50mL reagent reservoirs
- Seals for 96-well plates (e.g. Microseal B, BioRad MSB-1001)
- 96-well plate with flat transparent bottom for fluorometry (e.g. Corning 3603)
- SYBR Green I (Life technologies S-7563)
- DNA standards in range of 1-10ng/μl (we use those that come with Life Q-33120)
- Plate reader with SYBR-compatible filters

### Procedure

**Note:** This procedure assumes gDNA concentration in the range of 1-10ng/μl. If samples cover a different range of concentrations, the below procedure should be modified accordingly. We recommend a two step-dilution for samples with a broad range of concentrations.

1. Mix **5μl** of concentrated SYBR Green I with approximately **25mL** of TE in a clean reservoir to make a working dye solution.

- The same dye solution should be used across all samples and standards.
2. Seal, vortex and centrifuge gDNA (200rcf for 30s).
3. Add **10μl** of gDNA to each well of a fluorometry plate.

- Recommended: 96-well pipette or multi-channel pipette.
4. In 8 wells of another fluorometry plate, add **10μl** of each DNA standard.

- We found little plate-to-plate variability, as long as the same dye solution is used.
5. Dispense **190μl** of dye solution into each well.
6. Incubate the DNA and dye in the dark for 5 minutes, e.g., put in a dark drawer or covered with foil.
7. Read fluorescence on the plate reader using SYBR Green or GFP filters (e.g. Ex:470/40 and Em:520LP).

- If standards curve is not linear, allow the plate to sit longer and repeat.
- If the samples are outside the linear range of standards, repeat with a different volume of sample.
8. Based on measured DNA concentration of each sample, calculate the volume of water needed to dilute each sample to 0.5ng/μl.
9. Dispense these volumes of water into a fresh 96-well plate.

- Recommended: Liquid handling robot
- If doing manually, a tablet and Pippette-Guide-96 may be helpful (https://github.com/tamilieberman/Pipette-Guide-96).
10. Add **5μl** of gDNA to each well of the new plate.
11. Seal the plate tightly, vortex, and centrifuge (200rcf for 30s).

## Module 2. Tagmentation

**Goal:** Mix 1.25μl of TD buffer, 0.25μl TDE1, and 1μl of gDNA in each well. Final total volume per well is 2.5μl. Carry out tagmentation reaction in a thermocycler.

### Materials and equipment

- Normalized gDNA from Module 1
- Nextera TD buffer and TDE1 enzyme (from Illumina kits FC-121-1030 or FC-121-1031)
- 96-well PCR plate (e.g. Bio-Rad MLP-9601)
- Microseal ‘B’ (Bio-Rad MSB-1001)

### Procedure

**Note:** All reagents should be kept on ice. The 96-well plate containing samples should also be kept on ice while assembling the mix, and all steps should be done quickly.

1. Preheat thermocycler to 55°C. If starting from frozen gDNA, thaw, vortex and spin down gDNA. Thaw TD buffer and TDE1 on ice.
2. Invert TD buffer and TDE1 gently to mix, spin down, and replace on ice.
3. Make the tagmentation master mix (TMM) by mixing **156μl** buffer and **31.2μl** enzyme in a PCR tube. Mix thoroughly by gently pipetting up and down 10 times.

- Excess volumes enable rapid and even distribution of tagmentation enzyme and buffer. It is essential that all samples receive the same amount of enzyme.
4. Distribute TMM into 8-tube PCR strip with **22.5μl** per tube. Cap and spin down to remove bubbles.
5. Distribute **1.5μl** of TMM per well into all wells of a fresh PCR plate using a multi-channel pipette.

- Dispense into the bottom of the well and ensure the full amount is dispensed each time.
6. Transfer **1μl** of gDNA into each corresponding well using a manual multi-channel pipette. Mix by gently pipetting up and down 10 times.
7. Cover plate with Microseal ‘B’ and spin down (280rcf for 30s).
8. Incubate in thermocycler for **10min** at **55°C**.

- Incubation time between 5 and 20 minutes does not affect the results.
9. Place plate on ice. Allow plate to cool before proceeding to Module 3.

## Module 3. Adapter addition and library amplification

**Goal**: Mix 11μl of PCR master mix, 4.4μl of index1, 4.4μl of index2, and 2.5μl of tagmented DNA in each well. Final total volume per well is 22.5μl.

### Materials and equipment

- Tagmented DNA from Module 2
- KAPA 2X Library Amplification Kit (KAPA KK2611)
- Rxxx/Sxxx primers at concentration of 5μM, arrayed in PCR strip (8), supplemental file 1
- Cxxx/Nxxx primers at concentration of 5μM, arrayed in PCR strips (12), supplemental file 1
- Microseal ‘A’ (Bio-Rad MSA-5001)

### Procedure

**Note:** First 8 steps can be done during the tagmentation reaction (Module 2, step 8). Special care should be taken to avoid primer cross-contamination; only fresh tips should be inserted into the primer stock. It saves significant effort to aliquot the primers into 8- and 12-strip PCR tubes, so that multi-channel pipettes can be used.

1. Preparation. Thaw indexing primers at room temperature. Invert gently to mix. Spin down. Record which indexing primers you are using. Thaw the KAPA master mix at room temperature. Invert gently to mix.
2. Label one fresh 8-well PCR strip for the row master mix (RMM) and one fresh 12-well PCR strip for the column master mix (CMM).

- It is easy to accidentally rotate these strips—proper labeling is essential.
- Place RMM and Rxxx/Sxxx primer strips to the left of the PCR plate.
- Place CMM and Cxxx/Nxxx primer strips to the top of the PCR plate.
3. Distribute **85.8μl** of 2x KAPA master mix into each empty RMM tube.
4. Distribute **57.2μl** of 2x KAPA master mix into each empty CMM tube.
5. Add **68μl** of each Rxxx/Sxxx primer into the corresponding RMM tube. Mix by pipetting up and down, cap, and spin down.
6. Add **57.2μl** of each Cxxx/Nxxx primer into the corresponding CMM tube. Mix by pipetting up and down, cap, and spin down.
7. Transfer **10μl** of RMMs into each well of the tagmentation plate using a multi-channel pipette, so that each row receives the same Rxxx index.

- Change tips after every transfer.
- Make sure that the row number corresponds to the Rxxx.
8. Transfer **10μl** of CMMs into each well of the plate using a multi-channel pipette, so that each column receives the same Cxxx/Nxxx index. Mix by gently pipetting up and down 10 times.

- Change tips after every transfer.
- Make sure that the column number corresponds to the Cxxx.
9. Cover plate with Microseal ‘A’. Spin down (200rcf for 30s).

- Gently press the seal on each well, especially edge wells before spinning down; this seal is non-adhesive until heat is added.
10. Place plate in the thermocycler.

- Ensure that the lid is tight and is heated during thermocycling.
11. Run the following program:

1. 72°C for 3 min
2. 98°C for 5 min
3. 98°C for 10 sec
4. 62°C for 30 sec
5. 72°C for 30 sec
6. Repeat steps (3)-(5) 13 times for total of 13 cycles
7. 72°C for 5 min
8. Hold at 4°C

## Module 4. PCR clean-up and size selection

### Materials and equipment

- Tagmented and indexed DNA
- 100% ethanol
- Resuspension buffer (10mM Tris-Cl [pH 8.0] + 1mM EDTA [pH 8.0] + 0.05% Tween-20)
- Deep-well 96-well plate for bead purification
- Magnetic beads for DNA purification (e.g. according to http://ethanomics.files.wordpress.com/2012/08/serapure_v2-2.pdf)
- 96-well plate magnetic stand (e.g., Life Technologies, Cat. #123-31D).

### Procedure

**Note:** It is best to thaw beads at the beginning of the day, as it takes time for them to reach room temperature. While at this stage cross-contamination is a much smaller issue, we still recommend fresh tips for each well.

1. Centrifuge the PCR plate at 200rcf for 30 seconds.
2. To resuspend beads, alternate between vortexing and inverting beads for a total of at least 60 sec.
3. Transfer at least **2.5ml** of beads into a reagent reservoir.
4. Using a multi-channel pipette, transfer **22.5μl** of beads into each well of the PCR plate and pipette up and down several times to mix.

- Pipette into the bottom of wells and ensure complete dispense.
- Some prefer to purify in a separate deep-well 96-well plate, for easier aspiration. In this case, we recommend first adding 15μl of beads to each well of the deep-well plate, then transferring 15μl of sample into each well and mixing.
5. Incubate at room temperature for **5 min**. DNA is now on the beads.
6. Prepare a fresh batch of 80% ethanol by mixing **10mL** of sterile water and **40mL** of 100% ethanol in a sterile reservoir.
7. Place the plate on the magnetic stand and incubate for **1 min** to separate beads from solution. The solution should become clear.
8. While the plate is on the magnetic stand, aspirate clear solution from the plate and discard. Do not disturb the beads. If beads are accidentally aspirated, resuspend them, wait 1 min, and aspirate again.
9. While the plate is on the magnetic stand, dispense **200μl** of 80% ethanol into each well. Incubate for **1 min** at room temperature.
10. Aspirate ethanol and discard. Do not disturb the beads. If beads are accidentally aspirated, resuspend them, wait 1 min, and aspirate again.
11. Repeat steps 9-10 for a total of 2 washes.
12. Remove any visibly remaining ethanol droplets with smaller pipette tips.
13. Let the plate air dry for **20 min** for residual ethanol to evaporate.

- This is a good time to take out the gel-dye matrix for the BioAnalyzer in Module 5, to allow it to come to room temperature.
14. Transfer at least **3.5ml** of resuspension buffer to a new reservoir.
15. Take the plate off the magnetic stand. Add **30μl** of resuspension buffer to each well of the plate using a multichannel pipette. Resuspend the beads by mixing 10-15 times.
16. Incubate for **5 min** at room temperature. DNA is now in the solution.
17. Place the plate back onto the magnetic stand and incubate for about **1 min** to separate beads from solution. The solution should become clear.
18. While the plate is on the magnetic stand, aspirate clear solution from the plate and transfer to a fresh 96-well plate. Do not disturb the beads. If beads are accidentally pipetted, resuspend them, wait for the solution to become clear, and repeat.
19. Seal plate and spin down (200rcf for 30s).

## Module 5. Library QC and Pooling

### Materials and equipment

- Purified Nextera libraries from Module 4
- High Sensitivity DNA kit for BioAnalyzer (Agilent 5067-4626)
- TE buffer
- 50mL reagent reservoirs
- 96-well plate with flat transparent bottom for fluorometry (e.g. Corning 3603)
- SYBR Green I (Life technologies S-7563)
- DNA standards in range of 1–10ng/μl (we use those that come with Life Q-33120)
- Plate reader with SYBR-compatible filters
- BioAnalyzer (Agilent 2100) or similar DNA fragment-size assay system.

### Procedure

1. Perform steps 1-7 of Module 1 to quantify DNA concentration across all samples.

- You may use less DNA and ladder (5 or 8 μl) to conserve sample.
2. Calculate the concentration of each sample. Discard samples with concentration less than 0.5ng/μl.
3. Select a number of samples per batch (96-well plate) for fragment quantification.
4. Prepare High Sensitivity DNA Analyzer chip per manufacturers instructions, load samples onto chip, and perform analysis.
5. Determine if fragment-size distributions in each batch are acceptable. See Figures 2 and 3 for reference and main text for discussion. Calculate sample molarity based on batch-specific average fragment length.
6. Pool acceptable samples in equimolar concentrations.
7. If sequencing in-house, accurate quantification is crucial to achieve optimal cluster density. We recommend using qPCR and running a BioAnalyzer on the final pooled library.

## Acknowledgements

The authors thank Sheri Simmons, Evgeni Frenkel, and Lealia Xiong for help in developing this protocol. This work was supported by a National Science Foundation Mathematical Sciences Postdoctoral Research Fellowship (MB), a Career Award at the Scientific Interface from the Burroughs Wellcome Foundation (SK), by the James S. McDonnell Foundation, the Alfred P. Sloan Foundation, grant PHY 1313638 from the NSF, and grant GM104239 from the NIH (MMD) and by US National Institutes of Health grant R01-GM081617 (RK) and the European Research Council FP7 ERC Grant 281891 (RK) and Hoffman-LaRoche. Illumina requests the following statement and disclaimer to be added to publication of the alternate barcode sequences: “Oligonucleotide sequences © 2007–2011 Illumina, Inc. All rights reserved. Derivative works created by Illumina customers are authorized for use with Illumina instruments and products only. All other uses are strictly prohibited.”

